# The Influence of Brain MRI Defacing Algorithms on Brain-Age Predictions via 3D Convolutional Neural Networks

**DOI:** 10.1101/2023.04.28.538724

**Authors:** Ryan J. Cali, Ravi R. Bhatt, Sophia I. Thomopoulos, Shruti Gadewar, Iyad Ba Gari, Tamoghna Chattopadhyay, Neda Jahanshad, Paul M. Thompson, the Alzheimer’s Disease Neuroimaging Initiative

## Abstract

In brain imaging research, it is becoming standard practice to remove the face from the individual’s 3D structural MRI scan to ensure data privacy standards are met. Face removal - or ‘defacing’ - is being advocated for large, multi-site studies where data is transferred across geographically diverse sites. Several methods have been developed to limit the loss of important brain data by accurately and precisely removing non-brain facial tissue. At the same time, deep learning methods such as convolutional neural networks (CNNs) are increasingly being used in medical imaging research for diagnostic classification and prognosis in neurological diseases. These neural networks train predictive models based on patterns in large numbers of images. Because of this, defacing scans could remove informative data. Here, we evaluated 4 popular defacing methods to identify the effects of defacing on ‘brain age’ prediction – a common benchmarking task of predicting a subject’s chronological age from their 3D T1-weighted brain MRI. We compared brain-age calculations using defaced MRIs to those that were directly brain extracted, and those with both brain and face. Significant differences were present when comparing average per-subject error rates between algorithms in both the defaced brain data and the extracted facial tissue. Results also indicated brain age accuracy depends on defacing and the choice of algorithm. In a secondary analysis, we also examined how well comparable CNNs could predict chronological age from the facial region only (the extracted portion of the defaced image), as well as visualize areas of importance in facial tissue for predictive tasks using CNNs. We obtained better performance in age prediction when using the extracted face portion alone than images of the brain, suggesting the need for caution when defacing methods are used in medical image analysis.

## I. INTRODUCTION

Digital removal of the face from brain MRI scans has been advocated to safeguard privacy in neuroimaging research, but its effects on downstream algorithms have not been fully evaluated. Early work has proposed using manually-labeled non-brain facial features (mouth, ears, nose) to create a group template that can then be used as a mask to remove facial regions from newly acquired raw images. Probabilistic maps are then generated and any voxels within the image that have a nonzero chance of being brain tissue are retained [1]. As human facial features vary considerably across individuals, this form of defacing may not generalize well to thousands of datasets. Though these facial erosion techniques have been proposed, the use of defacing - and even small variations in the extent of defacing, such as removal of regions where there was signal dropout - may influence the accuracy of downstream analyses and research findings [2]. Prior work has assessed the impact of defacing algorithms on downstream analyses by examining the performance of skullstripping algorithms after defacing and found that defacing prior to skullstripping significantly improved skullstripping accuracy [1]. T1-weighted tissue segmentation accuracy has been assessed using a series of defacing algorithms with FreeSurfer’s ‘*recon-all’* pipeline and its tissue segmentation routine [3]. Similarly, a separate study used a tissue segmentation algorithm found in the Statistical Parametric Mapping (SPM) software suite, Computational Anatomy Toolbox (CAT) segmentation, and FreeSurfer’s recon-all and found that tissue segmentations only differed in one of the segmentation routines - CAT - depending on the defacing routine [4]. These performance assessments have measured accuracy in tissue segmentation, signal normalization, and percent overlap with the original image and mask image after defacing. Alternatively, image registration accuracy to the MNI-152 brain template has also been used to assess the influence of defacing algorithms [1]. One study examined performance based on the lasting presence of facial features and incorrect removal of brain tissue. It was found that across multiple datasets the defacing method included in the Analysis of Functional Neuro Images (AFNI) software suite was the most accurate, while the python-based tool, Pydeface had lower accuracy, depending on the dataset. Additionally, the age of the subject played a role in performance [2][5-6].

With the increased use of machine learning in medical imaging, it is important to retain as much data as possible to train and test a model. Removal of the face has been largely unexamined when using data-driven analytic models, such as those used in deep learning. Convolutional neural networks (CNNs), for example, are increasingly used for tasks such as brain age estimation - where an algorithm is trained to predict a person’s age from their MRI scan, after being trained on scans from healthy controls. Such brain age estimators have been used as a proxy for overall brain health, to identify lifestyle and genetic factors that promote or resist brain aging. People whose brain age is higher than their chronological age may be considered as having accelerated brain aging and perhaps being at risk for various age-related neurodegenerative diseases [7]. CNNs have also been applied to MRIs to classify individuals with degenerative diseases and predict future decline [7-8]. As CNNs apply kernels to images to compute feature maps, every voxel potentially holds informative data. However, no study, to our knowledge, has assessed whether facial features are contributing to brain age prediction.

Here we assessed the downstream effects of four popular defacing algorithms (AFNI, FSL, Pydeface, SPM) on the performance of age prediction models using a 3D-CNN. We did not use any data-driven deep learning algorithms for the defacing step, as we are currently unaware of any such methods, although they do exist for ‘skull stripping’ purposes. We also assess how well the face alone - without the skull or brain - could be used to predict biological age, as well as the predictive value of keeping the brain and face together. Predicting chronological age from 3D MRI data using an individual’s face may indicate whether facial features are strong contributors in predicting “brain-age”. By evaluating predictive performance when both brain and face are present in the image, we are able to indirectly identify which regions (either brain or face) are most predictive in age-related tasks. This is particularly relevant if defacing is conducted and pieces of facial tissue are left behind.

## II. METHODS

### A. Data Preprocessing

3D T1-weighted brain MRI scans of 1,343 cognitively healthy participants (age range: 56.1-95.8 years; 624 male, 718 female) from the Alzheimer’s Disease Neuroimaging Initiative phase 2 (ADNI2) dataset were used to assess brain-age prediction accuracy. To increase the training set size, subjects with multiple scans at different time points were used, along with data augmentation. All subjects underwent 4 different defacing protocols: (1) FSL [9], (2) SPM12 [10], (3) AFNI [5], and (4) a Python-based tool, “Pydeface” [6]. Only tools that are actively being used and maintained by their developers were evaluated. Anatomical locations of extracted facial tissue from each algorithm are highlighted in blue in **Figure 1**, for a typical healthy subject. AFNI removes large portions of the face, nose, jaw, and portions of the ears. FSL removes smaller portions of the face, minimal ear tissue, and the nose. Pydeface removes large portions of the neck, and the nose, but leaves the ears intact. SPM uses a 3D plane-like removal of the face and retains the ears. After defacing, all data underwent minimal preprocessing consisting of reorientation to a common standard space using FSL’s FLIRT linear registration tool (MNI-152), ANTS N4BiasFieldCorrection, brain extraction from the skull and surrounding tissue (HD-BET), linear registration to the UK Biobank standard template, and downsampling to a voxel dimension of 2×2×2 mm [11-14]. A skullstripped group that did not undergo defacing - but did undergo all other preprocessing steps - was used for comparison. Additionally, a dataset with the brain and face still intact (no skullstripping or defacing) was used to identify if and where predictive value remained when both brain and face were present. Lastly, we would like to note that per the requirement of FSL’s *deface*, standard space reorientation was applied before defacing in this specific case.

**Figure 1.**
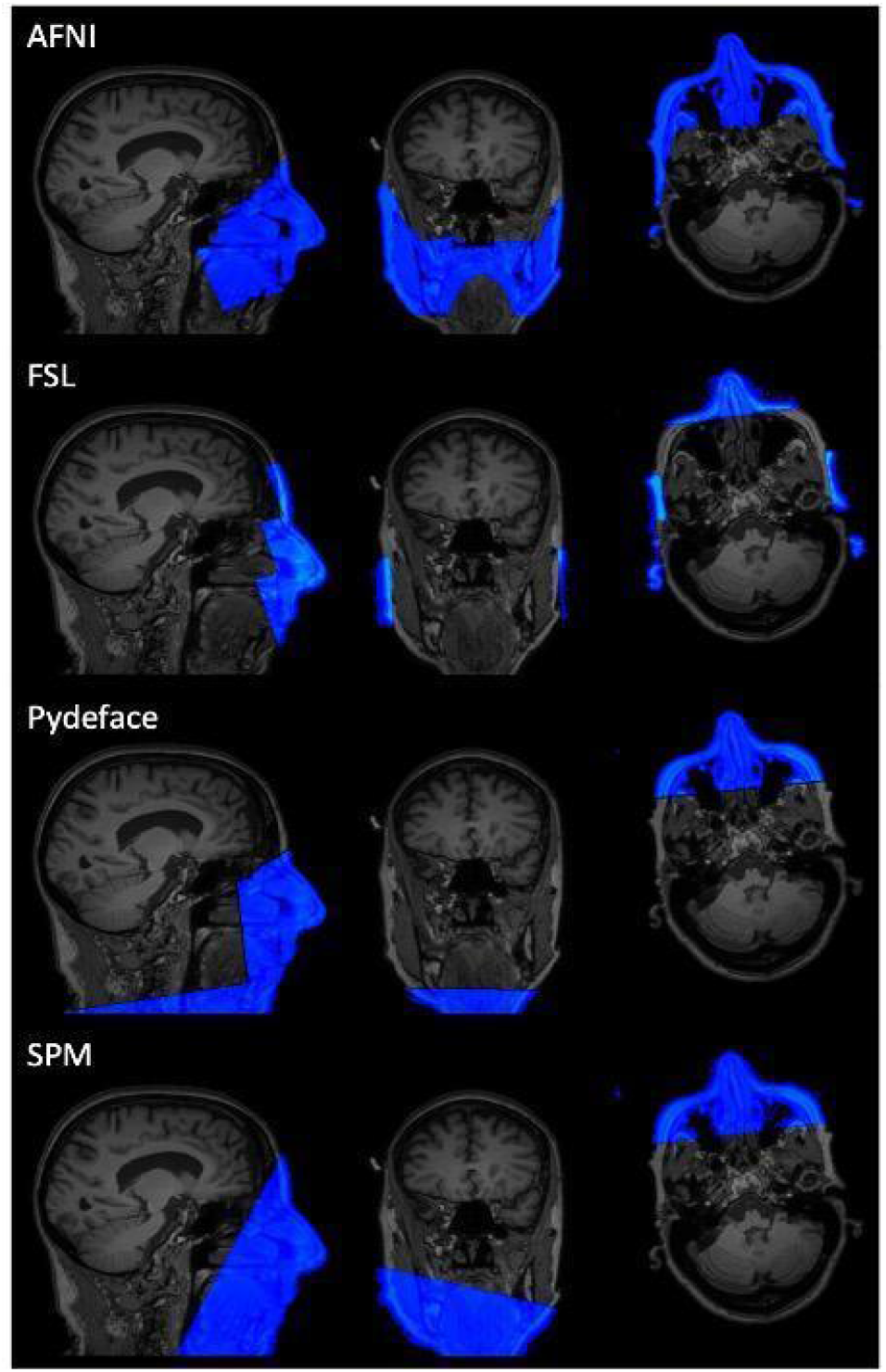
Extracted regions of non-brain facial tissue by each algorithm (*blue*).

### B. Convolutional Neural Network

For each defaced dataset, as well as the skullstripped and brain and face groups, data was split into the same 940 training, 267 validation, and 136 test samples. The input of the 3D CNN was an image with matrix dimensions of 91×109×91 and voxel dimensions of 2×2×2mm. The model consisted of three convolution blocks with filter sizes 128, 64, and 64 and additional layers of instance normalization and max pooling, and was trained for 100 epochs, with a learning rate of 1e-4 and batch size of 4 to reduce computational cost on our computing cluster (**Figure 2**). To increase the size of the training data, we used data augmentation in the form of a set of random rotations. The initial learning rate was 1e-4 and was exponentially decayed with a decay rate of 1e-4. The Adam optimizer and MSE loss function were used for training. Early stopping was used to avoid overfitting of the data. Each model was run 3 times to generate an average mean absolute error (MAE) estimate across runs. There were a total of 821,377 trainable parameters. The network architecture is shown in **Figure 2**.

**Figure 2.**
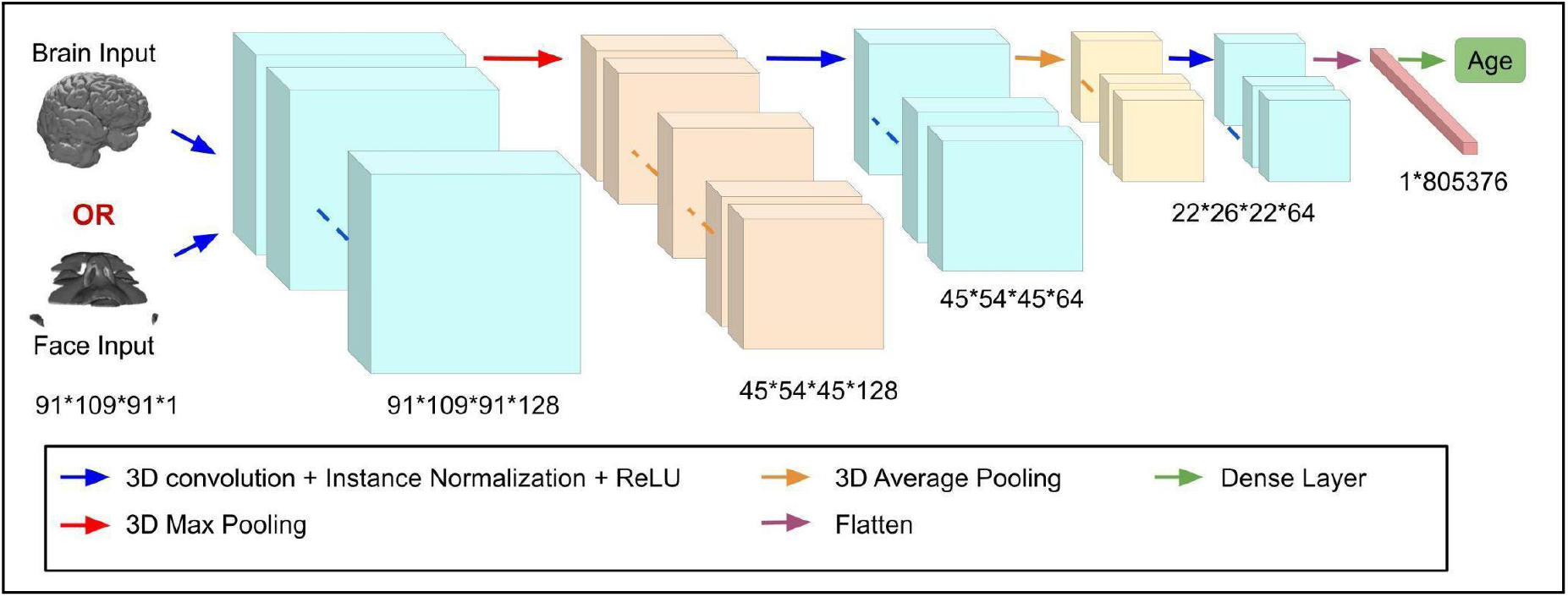
Visual representation of the 3D CNN model used for brain-age prediction.

### C. Saliency Maps

Saliency maps allow the interpretation of hidden layers in deep neural networks using gradients of the output over the input layer. This can be useful when attempting to understand how a given model “sees” the input data as it passes through the various layers of the network. Specifically, visualization of ‘heat maps’ (red-yellow indicating greatest predictive significance and green-blue indicating lowest) enables us to see areas with the greatest importance for predictive models in image classification. Here, we employed the Gradient-weighted Class Activation Mapping (Grad-CAM) technique to visualize the areas of most predictive importance in the images in the final convolutional layer of the model [15]. This technique has been used extensively on 3D brain images to identify cortical regions of most importance for a given predictive task [16]. However, to our knowledge, this method has not been used on extracted facial tissue, or intact brain scans with the face still present in the image. Saliency maps were generated for both the facial tissue (blue regions in **Figure 1)**, as well as the brain and face images using the Grad-Cam tool within the Keras API. The resulting maps - from one randomly selected individual - show spots of greater predictive value in the jaw region of the extracted facial tissue shown in the sagittal plane in **Figure 3**. In the brain and face images, the face was consistent in being the area with the greatest predictive value (**Figure 3**).

**Figure 3.**
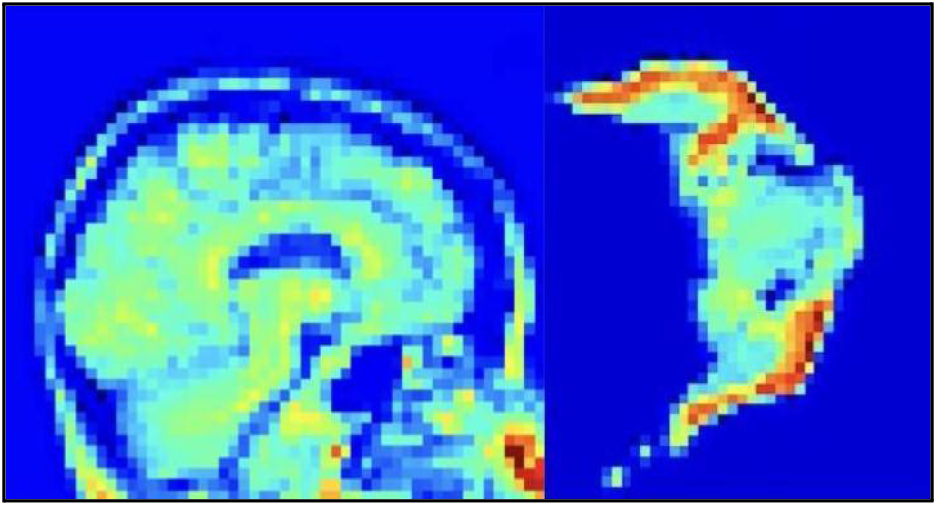
Heatmap showing a visual representation of the final convolutional layer in the CNN when predicting age. As noted by the red colors in the highlighted areas, at least some facial regions contribute to the brain-age prediction, in line with our findings that deleting them from the image affects algorithm performance. Pictured: Sagittal view of the skull and brain (left); Overhead view of the face (right).

**Figure 4.**
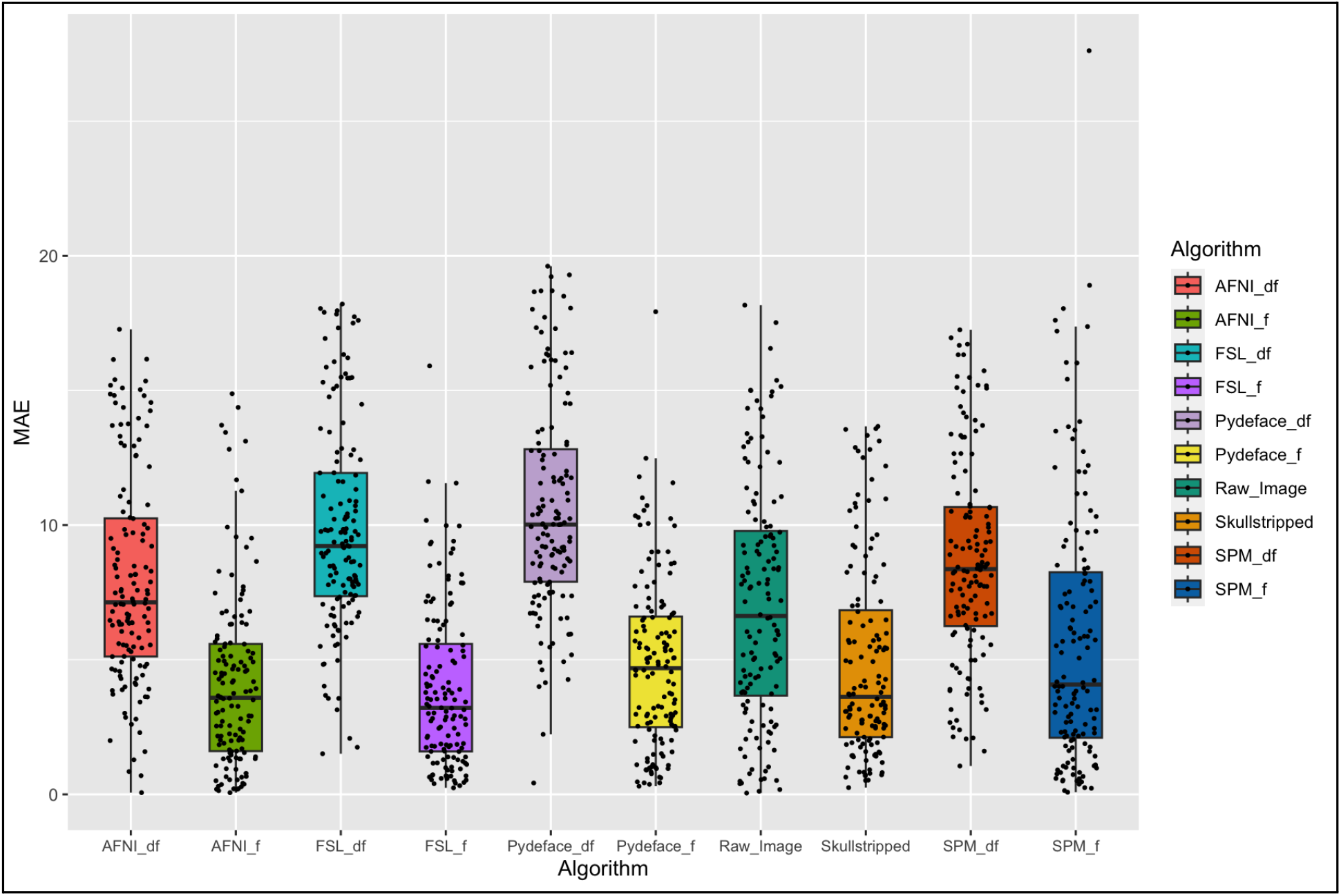
Graph showing per-subject distributions of MAE values in the test set (N = 136). *X*-axis labels with ‘df’ denote defaced data; ‘f’ facial tissue data (i.e., the extracted facial portion from each image, only)..

## III. RESULTS

The mean absolute error (MAE) was used as a performance metric for all brain age calculations, and the MAEs for all models are shown in **Table 1**. For every defacing method tested, the MAE was higher after defacing the scans, indicating poorer performance.

**TABLE 1.** Mean Absolute Error (MAE) estimates in Years.

| Experiment | MAE |  |  |  |
| --- | --- | --- | --- | --- |
|  | Run 1 | Run 2 | Run 3 | Mean |
| **AFNI Defaced** | 7.75 | 8.07 | 8.07 | **7.96**<br>(3.93) |
| **AFNI Face-only** | 4.17 | 4.13 | 4.13 | 4.14<br>(3.27) |
| **FSL Defaced** | 8.55 | 12.49 | 8.48 | 9.84<br>(3.86) |
| **FSL Face-only** | 3.92 | 4.05 | 3.73 | **3.90**<br>(2.97) |
| **Pydeface Defaced** | 10.90 | 10.69 | 10.29 | 10.63<br>(4.00) |
| **Pydeface Face-only** | 5.36 | 3.99 | 5.13 | 4.83<br>(3.18) |
| **SPM Defaced** | 8.74 | 8.28 | 9.12 | 8.71<br>(3.85) |
| **SPM Face-only** | 5.74 | 5.84 | 5.84 | 5.8<br>(4.99) |
| **Skullstripped (no defacing)** | 4.91 | 5.37 | 4.39 | **4.89**<br>(3.62) |
| **Raw Image (Brain and Face)** | 7.06 | 7.06 | 7.06 | **7.06**<br>(4.03) |

The defacing method within Analysis of Functional Neuroimages toolbox (AFNI) performed the best (MAE: 7.96) and Pydeface performed the worst (MAE: 10.63). The skullstripped dataset that only underwent brain extraction and no defacing achieved an MAE of 4.89 and the brain and face data that did not undergo defacing or skullstripping, an MAE of 7.06. In the data that only contained extracted facial regions, the facial tissue removed by FSL’s defacing algorithm gave the best performance in estimating age in controls (MAE: 3.90); performance was the poorest when using SPM (MAE: 5.8).

A one-way ANOVA was performed to compare the effect of defacing algorithm on average error rate in brain-age and face-age prediction in the 136-subject test set. Significant differences were present between modalities *F*_*(9,1350)*_ = 53.83, p<0.0001. Tukey’s honest significance test revealed significant differences in defacing algorithms (p<0.05; CI = 95%) between SPM and Pydeface (p = 0.001, [0.431,3.396]), SPM and skullstripped (p <0.0001, [-5.308,-2.344]), AFNI and FSL (p = 0.0025, [-3.359,-0.394]), AFNI and Pydeface (p < 0.0001, [-4.146,-1.182]), AFNI and skullstripped (p < 0.0001, [1.594,5,4.558]). For the data that included the brain and face, significant differences were present when comparing with defaced SPM data (p = 0.0155, [-3.133,-0.169]), defaced FSL (p < 0.0001[1.296,4.260]), and defaced Pydeface (p < 0.0001, [2.083,5.047]). When comparing the extracted facial tissue from each algorithm, significance was found between SPM and AFNI (p = 0.0145, [-3.142,-0.178]), SPM and FSL (p = 0.0021, [-3.383,-0.4188]). Brain and face data showed significant differences with facial tissue when compared with FSL (p < 0.0001, [-4.643,-1.679]), and Pydeface (p < 0.0001, [-3.718,-0.753]) and skullstripped data (p = 0.0002, [0.6928,3.657]).

## IV. DISCUSSION

In this study, we evaluated the downstream impact of several popular defacing algorithms used to de-identify brain images. We show that specific algorithms perform better than others at retaining tissue of importance for accurately predicting brain age. Most notably, by performing skull stripping only (removing all non-brain tissue including the scalp) without running any specific defacing algorithm beforehand, mean absolute error (MAE) was lowest (better). The Python-based tool, “Pydeface”, generated the highest MAE score (poorer). This might persuade one to use skull stripping rather than defacing, but various multisite data-sharing studies may require data to be defaced prior to sharing with outside collaborators, adding an additional processing step that would need to be harmonized.

An additional experiment was conducted using only the extracted facial portions, as well as the brain and face together (‘raw image’) to predict chronological age. Facial tissue was sufficient to give good performance in age prediction tasks, with performance being better than when using the extracted brain. In the brain and face dataset, performance was poorer than just the face alone and greater than when defacing was conducted (MAE: 7.06). Much prior work has aimed to predict a person’s chronological age from 2-dimensional photos using CNNs [17] although we do not know of prior work predicting age from facial regions of brain MRI.

Performance on the facial tissue-based -age prediction task was best for FSL’s defacing routine. As shown in **Figure 1**, FSL tends to remove more of the “identifiable” portions of the image (face, ears, nose, mouth) than the 3 other approaches, which could explain the high predictive accuracy on the downstream task (MAE: 3.90). Conversely, SPM uses a plane-like extraction of facial tissue and gives poorer downstream accuracy (MAE: 5.8) than the skullstripped-only dataset (MAE: 4.89). However, unlike the other routines, FSL required the data be reoriented to standard space prior to defacing, which may explain some of the difference in performance.

As one limitation of the current approach, it could be that the face may not be as relevant for deep learning tasks that have to do with brain disease – such as classification or prognosis of Alzheimer’s or Parkinson’s disease. Arguably, a brain disease detection method should not need to use the face at all - but the presence of correlations and mutual information between the facial signals and the rest of the image means that we cannot rule out that the face contributes useful data. We chose the current task of brain age estimation as the ground truth is known for this task. Our results show that, in practice, deep learning algorithms are not invariant to face removal, unless they are restricted to learning features from a brain mask only (which, as shown here, may not be optimal). Future work will focus on other disease-relevant tasks.

We hope that this work will offer insights for those analyzing shared multi-site data on which methods preserve accuracy while minimizing security concerns for patients and research volunteers. We did not address whether the algorithms do indeed safeguard privacy, and they may not, given work on generative adversarial methods that aim to put the face back on defaced images [18-19]. Defacing effects were only tested here with one specific brain-age prediction model on a moderate-sized dataset, and our work will benefit from assessing results from other tasks, including diagnosis and segmentation, as well as using larger sample sizes.

Outside of defacing, work is currently being conducted to introduce means of “refacing” neuroimaging data in order to mitigate membership attacks (re-identifying an individual) through data breaches. In some approaches, a population average face is merged with an existing image to remove a given individual’s identifying facial features. This work proved to be about as useful in preventing such membership attacks as the FSL-based defacing algorithm [20]. Though promising, based on our findings presented here, great care must be taken to ensure that the data is not manipulated in such a way that discards useful data for downstream analyses.

## V. CONCLUSION

A key result of this study is that defacing algorithms interfered with the performance of all the deep learning methods that we studied, on the task of estimating a person’s age from their MRI. After defacing, with all methods, the mean absolute error for brain age estimation was higher than without defacing. Facial features improve performance on this task. Here we show that brain-age and face-age are two distinct outputs and that any facial tissue that is potentially left behind after defacing could influence brain-age prediction performance. This work also highlights that there can be substantial variation in downstream results depending on whether the images are defaced, and which defacing algorithm is used.

## VI. ACKNOWLEDGMENTS

This work was supported in part by NIH grant U01AG024904 to the ADNI MRI Core, R01AG058854, R01AG059874, NSF 4AD Initiative. Data collection and sharing for the Alzheimer’s Disease Neuroimaging Initiative (ADNI) (National Institutes of Health Grant U01 AG024904) and DOD ADNI (Department of Defense award number W81XWH-12-2-0012). ADNI is funded by the National Institute on Aging, the National Institute of Biomedical Imaging and Bioengineering, and a variety of private funders, listed in full at the ADNI website: (https://adni.loni.usc.edu/wp-content/uploads/how_to_apply/ADNI_Acknowledgement_List.pdf).

